# Medial prefrontal cortex and nucleus reuniens are critical for working memory in an operant delayed nonmatch task

**DOI:** 10.1101/2024.07.03.601891

**Authors:** Evan J. Ciacciarelli, Scott D. Dunn, Taqdees Gohar, T. Joseph Sloand, Mark Niedringhaus, Elizabeth A. West

## Abstract

Working memory refers to the temporary retention of a small amount of information used in the execution of a cognitive task. The prefrontal cortex and its connections with thalamic subregions are thought to mediate specific aspects of working memory, including engaging with the hippocampus to mediate memory retrieval. We used an operant delayed-non match to position task, which does not require the hippocampus, to determine roles of the rodent medial prefrontal cortex (mPFC), the nucleus reuniens thalamic region (RE), and their connection. We found that transient inactivation of the mPFC and RE using the GABA-A agonist muscimol led to a delay-independent reduction in behavioral performance in the delayed non-match to position paradigm. Critically, we used a chemogenetic approach to determine the directionality of the necessary circuitry for behavioral performance reliant on working memory. Specifically, when we targeted mPFC neurons that project to the RE (mPFC-RE) we found a delay-independent reduction in the delayed non-match to position task, but not when we targeted RE neurons that project to the mPFC (RE-mPFC). Our results suggest a broader role for the mPFC-RE circuit in mediating working memory beyond the connection with the hippocampus.

## 1. Introduction

Working memory refers to the *online* retention and manipulation of a small amount of information to guide behavior trial by trial during cognitive tasks (Baddeley 1992). Impairments in working memory are critical features in both neuropsychiatric (e.g., schizophrenia) and neurodegenerative (e.g., Alzheimer’s disease, AD) disorders. Working memory performance depends on the prefrontal cortex (PFC) function. For example, patients with schizophrenia show disrupted prefrontal cortical activity linked to working memory deficits (Cho et al. 2006; Haenschel et al. 2009; Chen et al. 2014). In addition, PFC dysfunction (Salat et al. 2001; Kumar et al. 2017; Guarino et al. 2018) contributes to working memory deficits observed in people with mild cognitive impairment and AD (Kirova et al. 2015). While rats lack a direct, anatomical homolog to the PFC, there is a functional homology between the PFC in primates and the rodent medial prefrontal cortex; mPFC (Vertes 2004), including a role in working memory (Churchwell et al. 2010; Euston et al. 2012; Benoit et al. 2020).

The ability to hold information online and cognitively adjust behavior accordingly, a critical component of working memory, is controlled by the PFC, and the mPFC is critical for maintaining working memory across a wide range of demands. Specific roles of the mPFC have been found in spatial working memory (Urban et al. 2014), sequence memory (Jayachandran et al. 2019), and non-spatial working memory tasks (Kesner et al. 1996) suggesting a broad role in managing information online to guide behavior. In rodents, working memory is often determined in a wide range of tasks that also require spatial or temporal components that depend on the integrity of the hippocampus (Hampson et al. 1999; Nguyen et al. 2000; Zhang et al. 2013). However, as the mPFC does not project to the hippocampus directly (Sesack et al. 1989; Buchanan et al. 1994), the thalamic nucleus, the nucleus reuniens (RE), is thought to play a key role in mediating prefrontal-hippocampal interactions to guide the ability to retain information and use it to guide working memory (Vertes 2002; Dolleman-van der Weel et al. 2019). The hippocampus is also critical for delayed (non) matching performance in tasks that require spatial navigation in a T-maze or water maze (Hampson *et al*. 1999; Nguyen *et al*. 2000; Zhang *et al*. 2013), while the hippocampus is not necessary for performance in the *operant* version of the delayed (non)match to position task (Sloan et al. 2006). Thus, we aimed to determine if the circuitry between the mPFC and the RE is critical in the delayed non-match to position operant task to identify a potential role of these nuclei beyond hippocampal-dependent working memory.

Finally, we recently found transient differences in working memory between the Fischer 344 and the Long-Evans rats (Gohar et al. 2023). Thus, here we determined the role of the temporary inactivation of mPFC and RE circuitry in an operant delayed nonmatch to memory in Fischer 344 rats. Despite being frequently used in aging research (Beas et al. 2013; McQuail et al. 2016), the Fischer 344 strain is less often used in tasks assessing the neural circuits mediating cognitive function. In addition, Fischer 344 rats are the least similar in terms of their morphology, sensory and motor abilities compared to other standard laboratory rat strains (e.g. Lewis, Long-Evans, Sprague–Dawley, and Wistar rats) commonly used in assessing the circuits underlying working memory (Webb et al. 2003). Finally, the Fischer 344 strain is the background strain of a transgenic rat model developed to recapitulate age-dependent increases in AD pathology and memory deficits (Cohen et al. 2013; Rorabaugh et al. 2017). Given that working memory impairments are a common feature of AD (Ferretti et al. 2018; Mielke 2018), it is important to dissect the circuits critical for working memory in this strain (F344) to facilitate future study.

## 2. Materials and Methods

### 2.1. Subjects

In this study, we used male (n=19) and female (n=13) Fischer 344 rats aged 70-100 days (180- 270 grams for the females and 260-370 grams for the males) at the beginning of the study. Rats were acquired from Envigo (Livermore, California) or through our in-house breeding colony. All subjects were housed individually post-surgery and maintained on a reverse 12:12 hour light-dark cycle (lights off at 9:00 am). During behavioral training and testing, rats were restricted to no less than 85% of their body weight by food access to 10-25 g (5-6g/100g body weight) of standard rat chow per day (*ad libitum* water). All animal procedures were approved by the Rowan University Institutional Animal Care and Use Committee (IACUC).

#### Surgical Procedures: Viral Injection

For all experimental groups, 0.85 μl of virus coding for either muscarinic-receptor-based DREADD AAV8-hSyn-hM4Di(Gi)-mCherry or control virus AAV8-hSyn-EYFP (AddGene, Cambridge, MA) was injected bilaterally into either the mPFC (AP: + 2.7, ML: +/- 0.6, DV: -4.0 from skull) or the RE (AP: -2.5, ML: +/- 1.5, DV: -7.4 from skull, 10-degree angle) through a 30-gauge needle based on Paxinos and Watson standard rat brain atlas (Paxinos and Watson 2013). While we aimed our injection site at the prelimbic subregion within the mPFC, viral injections often spread beyond the prelimbic into the nearby infralimbic region, thus the entirety of the mPFC was likely affected, and both subregions project to the thalamic RE (Vertes 2002). Virus was injected sequentially into each hemisphere at a rate of 0.2 μl/min; the injection needle was left in place for at least 5 min to allow virus diffusion before retraction. Viral titers were > 7 x10^12^ vg/ml Following surgery, rats were given meloxicam (1 mg/kg, s.c.) for 3 days. After a week of recovery, rats were food regulated and begun training procedures.

#### Behavioral Training

We trained rats to perform a delayed nonmatch to position (DNMTP) task according to the timeline represented in Figure 1A. The training was performed in standard Med Associates operant chambers, consisting of two cue lights, two levers below the lights, a food receptacle between the levers for reward (sucrose) pellet delivery, a nose poke receptacle on the wall opposite to levers, and a house light. First rats underwent magazine training, in which a dustless precision sucrose pellet (Bioserve, Flemington, NJ, males 45 mg and females 20 mg) was delivered into the food receptacle pseudorandomly 30 times. After three days of magazine training, the rats were trained to press a lever on a fixed ratio schedule (FR), in which each lever press results in reward delivery. FR1 training consisted of four 15-minute blocks (2 minutes in between blocks). The rats were required to press each lever 50 times during this session to move to the next training stage. Next, rats were trained to self-initiate discrete trials. Here, each trial began with both levers retracted and the house light illuminated. For the levers to be extended into the chamber, the rat had to nose poke at the receptacle in the wall opposite to the levers within 20 seconds. After the nose poke, the house light turned off and one of the two cue lights turned on, and the lever below it was extended into the chamber. One press of the lever (FR1) was followed by a reward pellet delivery within 20 seconds of lever extension. The cue/lever position, left or right, was presented pseudorandomly so that each lever was presented 25 times (50 trials total). Failure to nose poke within 30 s of trial start resulted in the termination of that trial (omission) and the start of the next trial. To advance to the next stage of training, the rats were required to complete 45 trials out of 50 total possible trials (≥90%). The intertrial interval was pseudorandom and variable (85, 90, 95 s). After rats learned to self-initiate trials, we began delayed non-match to position (DNMTP) training (Gohar *et al*. 2023). First, each trial consisted of three phases i.e., sample phase, delay phase, and choice phase as shown in Figure 1B. Each session of this training stage consisted of 50 trials. During the *sample* phase, the house light was illuminated, and one of the two levers (e.g., right) located below it extended into the operant chamber (left or right position presented pseudorandomly). The rats were required to press the lever for a sucrose pellet on the sample phase to initiate the delay phase. During the delay, all lights were extinguished, and levers were retracted. After the delay, the house light was illuminated, and the rat had to nose poke to initiate the choice phase. During the *choice* phase, both levers extended into the chamber and the rat was required to press the opposite (i.e., non-match) lever from the sample phase (e.g., left) to obtain a sucrose pellet. Failure to press the sample lever, to nose poke during the delay phase, or to press a lever during the choice phase resulted in the termination of that trial, recording it as an omission. First, rats were trained with “no delay”; i.e., 1s between sample press and delay phase termination. After the advancement criterion was passed with no delay (≥88% correct), the rat was moved to the next stage of training which consisted of 50 trials on the same task except the delay phase was 2 seconds. After the advancement criterion was met (≥84 % correct), the rat was moved to the next stage of training which consisted of 50 trials with a 4-second delay. After reaching this criterion, rats underwent a second surgery for cannulae implants (see below). After 1 week of recovery, rats were trained one day with a 6-second delay. After training on the 6-second delay, rats were moved to the DNMTP multi-delay sessions. The rats were then moved through three multi-delay stages for 1 session each until testing began, stage 1 consisted of trials with short delay lengths (2, 4, 8, or 16 seconds), stage 2 consisted of medium delay lengths (4, 6, 12, and 24 seconds) and stage 3 consisted of trials with long delay lengths (4, 8, 16, 32 seconds). The delay periods were intermixed pseudorandomly with ∼15 trials/delay lengths for a total of 60 trials per session. Each trial in the multi-delay stages consisted of 3 phases, the sample, delay, and choice phase (Figure 1B). Test sessions (i.e., sessions in which the animal received an intracerebral injection before DNMTP) consisted of the delays in stage 3 (4, 8, 16, 32 seconds) as shown in Figure 1A-B. At least one baseline session in which no drug was delivered was run prior to each test session to ensure intracerebral injections did not interfere with performance post-test sessions and consisted of the same delays at in stage 3 (4, 8, 16, 32 seconds) as shown in Figure 1A-B.

**Figure 1.**
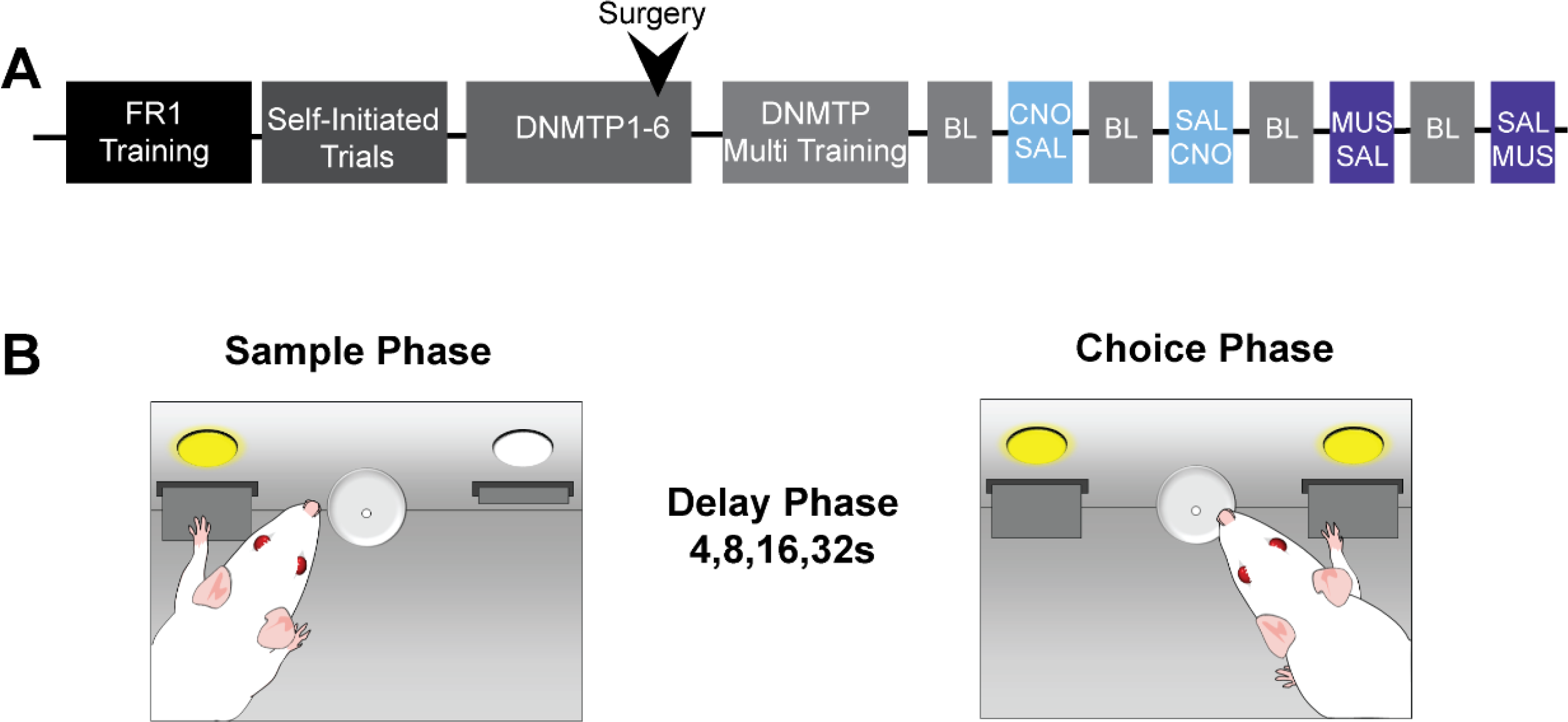
Schematic of the behavioral timeline and testing procedures. A) The timeline of behavioral training following virus surgery (FR-Fixed Ratio, DNMTP-Delayed Non-Match to Position with 1-4s delays) and after cannula surgery (DNMTP with multiple delays) with gray bars representing training including baseline (BL) sessions, light blue indicating either Clozapine-N-Oxide (CNO) or saline (SAL) and dark blue representing muscimol (MUS) or saline (SAL) delivered before the test ressions. B) A schematic of the DNMTP task with multiple delays (4, 8, 16, or 32s) used for training, baseline and test sessions representing three phases, sample, delay or choice.

#### Surgical Procedures: Cannulae Implant

Rats were implanted with guide cannulae (22 gauge; P1 Technologies, Roanoke, VA) fitted with 28-gauge internal cannulae that extended 1 mm beyond the tip of the guide in either the PrL (AP: 2.8, ML: +/- 1.5, DV: -4.2 from skull, 15-degree angle) or the RE (AP: -2.3, ML: +/- 2.4, DV: -7.8 from skull, 15-degree angle) and fixed to skull by screws using dental acrylic (West et al. 2012). Twenty-eight-gauge dummy cannulae (P1 Technologies, Roanoke, VA) cut to the same length or shorter as the guide cannulae were inserted to maintain cannula patency. Following surgery, rats were given meloxicam (1 mg/kg, s.c.) for 3 days. Rats recovered for one week before resuming behavioral training and testing (see above).

#### Drugs

Muscimol (R&D Systems, Minneapolis, MN) was dissolved in sterile saline to make a 500 nM solution. In the group of rats with cannulae aimed at the RE, a volume of 0.1 µl was infused into RE in each hemisphere. In the group of rats with cannulae aimed at the mPFC, a volume of 0.3 µl was infused into the mPFC in each hemisphere. Clozapine-N-oxide (CNO) hydrochloride (ChemSpecial, NC) was dissolved in sterile saline to make a 2.5 mM solution of CNO (Wicker et al. 2022). A volume of 0.3 µl was infused into the either the RE or mPFC in each hemisphere.

#### Intracerebral infusions

Internal infusion cannulae were attached to 10.0 µl Hamilton syringes via polyethylene tubing, which was filled with saline. A small air bubble separated the saline from the drug or vehicle. Drugs were infused bilaterally into mPFC or RE at a rate of 0.1 µl/min using an infusion pump (New Era Pumps Systems Inc, Model: 300, Farmingdale, NY), 5 minutes before behavioral testing. Rats received 4 intracerebral injections across the experiment (Figure 1A). On the first week of testing, rats received CNO or SAL, counterbalanced across animals. The subsequent week, rats received MUS or SAL, counterbalanced across animals. At least one baseline DNMTP training session was performed between intracerebral infusions test days as shown in Figure 1A.

### 2.4. Data Analysis

For pharmacological inactivation experiments, we recorded the % correct for each delay on the DNMTP task and ran a two-way repeated measures (RM) ANOVA with drug treatment (MUS vs Saline) and delay length as a repeated measures factor. We also analyzed these data with the average % correct across the delays and ran a paired student’s t-test (MUS vs SAL). For chemogenetic experiments, we recorded the % correct for each delay on the DNMTP task and ran a three-way mixed ANOVA with between-subject factors, virus (control vs hM4di) and drug treatment (CNO vs Saline), and delay length as a repeated measures factor. We also analyzed these data with the average % across the delays and ran a two-way mixed ANOVA with virus (control vs hM4di) as a between-subject factor and drug treatment (CNO vs SAL) as a repeated measures factor.

## 3. Results

### Pharmacological inactivation of the mPFC and RE decreases performance in DNMTP

We found that pharmacological inactivation in either the mPFC or RE by the GABA-A agonist, MUS, led to a delay-independent decrease in performance in the DNMTP. Specifically, a two-way RM ANOVA with Geisser-Greenhouse correction revealed a significant main effect of delay (mPFC: F (2.644, 47.60) = 17.42, p <0.0001; RE: F (2.620, 20.96) = 13.02, P<0.0001) and main effect of drug treatment (mPFC: F (1.000, 18.00) = 13.37, p=0.002; RE: F (1.000, 8.000) = 8.639 P=0.0187), but no interaction of delay and drug (mPFC: F (2.847, 51.25) = 0.1616; RE: F (2.899, 23.19) = 2.447, p=0.091) on the percent correct as shown in Figure 3. Since there was no interaction between drug and delay, we averaged the percent correct across all delays within each treatment condition to examine how MUS affected performance across individual rats. We found that when given MUS into the mPFC or RE rats show fewer correct responses as revealed by a paired students t-test (PrL: t=3.45, df=18, p=0.0026; RE: t=2.94, df=8, p=0.019).

**Figure 2.**
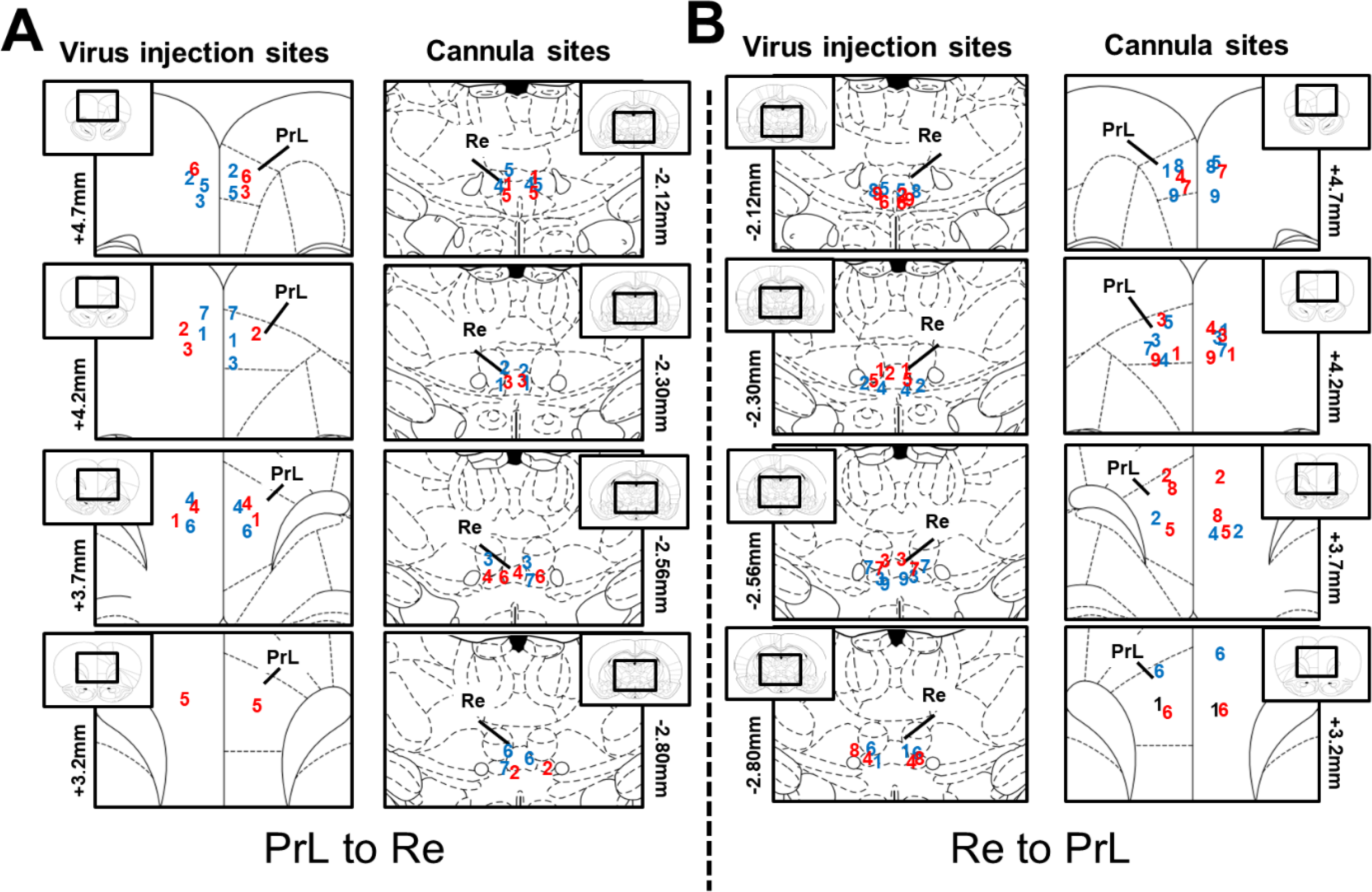
Viral and cannula placement for all rats in the mPFC-RE and RE-mPFC groups. A) Placement for viral injection sites and cannulas for rats the received virus (GFP, blue or hM4Di, red) in the medial prefrontal cortex (mPFC) and guide cannula in the nucleus reuniens (RE). B) Placement for viral injection sites and cannulas (GFP, blue or hM4Di, red) in the RE and guide cannula in the mPFC

**Figure 3.**
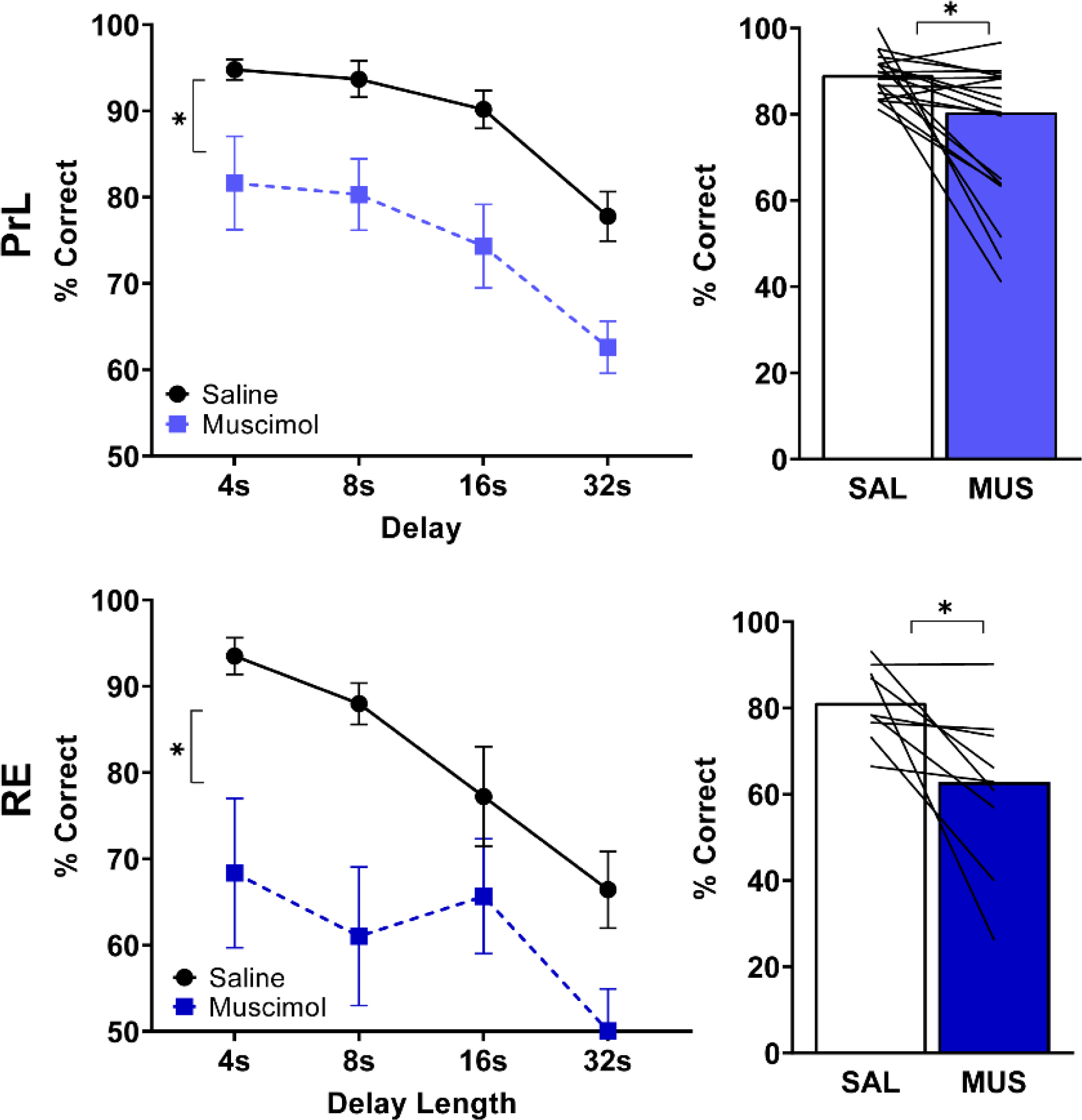
Pharmacological inactivation of the medial prefrontal cortex (mPFC) or nucleus reuniens (RE) impairs performance in an operant delayed nonmatch to position task. A) Pharmacological inactivation by the GABA-A agonist, muscimol (MUS), in the PrL leads to an overall decrease in the % correct in delayed non-match to position task across delays (* p<0.05 Saline vs MUS) B) individual rats overall percent correct averaged across delays (* p <0.05) shown across the session. C) Pharmacological inactivation in the RE leads to overall decrease in the % correct in the delayed non-match to position task across delays D) individual rats overall percent correct averaged across delays shown across the session (* p <0.05).

**Figure 4.**
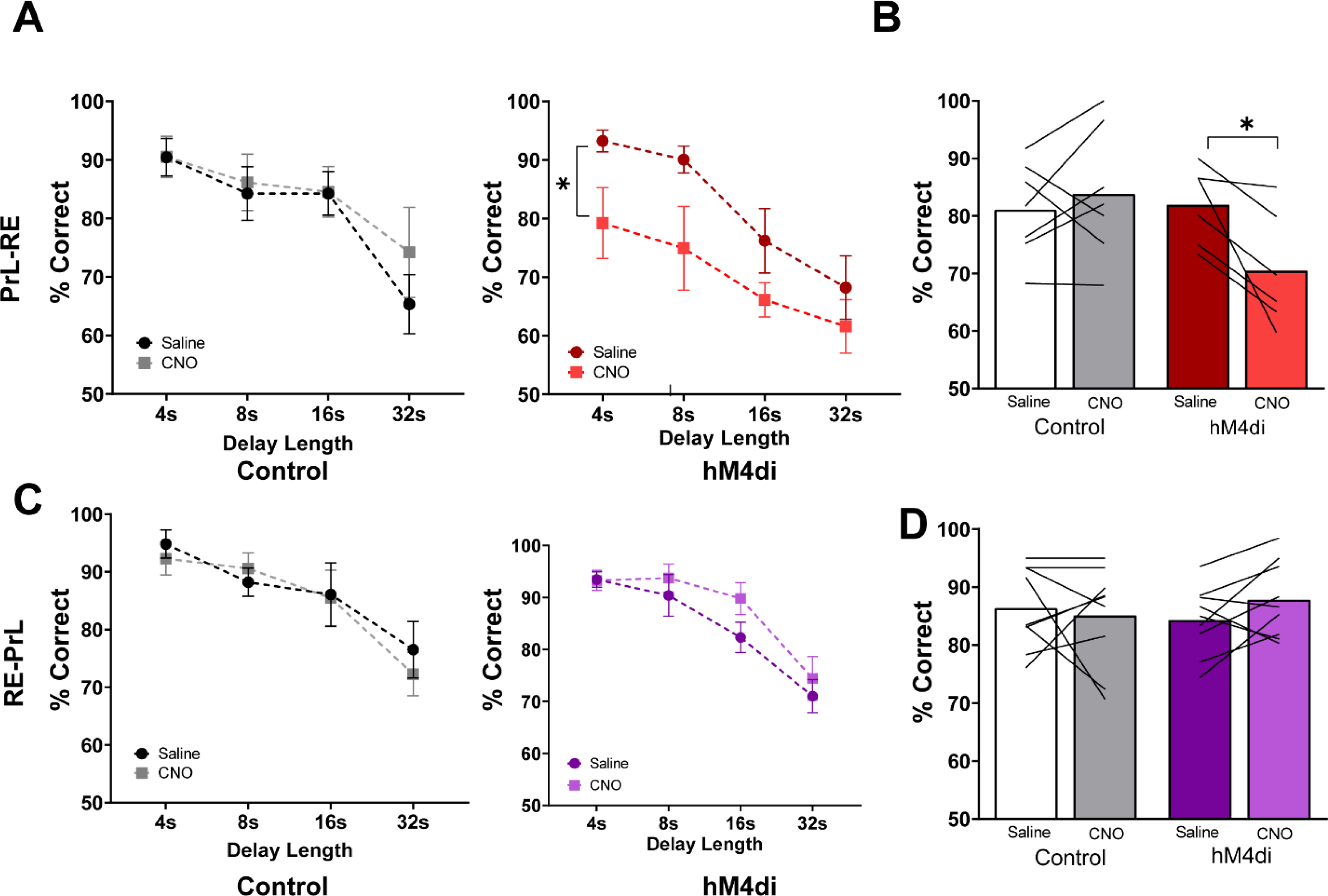
PrL-Re circuit, but not the Re-PrL circuity is critical for working memory in an operant delayed non-match to position task. A) Chemogenetic inhibition of the mPFC à RE circuit lead to an overall decrease in the % correct in delayed non-match to position task across delays (drug treatment x viral injection interaction effect, p<0.05, * CNO vs SAL post-hoc <0.05) B) individual rats overall percent correct averaged across delays (* p <0.05) shown across the session. C) Chemogenetic inhibition of the RE à mPFC circuit was without effect delayed non-match to position task across delays D) individual rats overall percent correct averaged across delays shown across the session.

### Silencing of mPFC-Re circuit, but not the Re-mPFC circuit, decreases performance in DNMTP

We found that using an inhibitory chemogenetic approach targeting PrL neurons that project to the RE led to a delay-independent decrease in performance in the DNMTP. We found that the average percent correct was decreased when rats received CNO in RE in rats that received DREADD virus hM4Di, but not in rats that received the control virus in the PrL. Specifically, a three-way mixed ANOVA with Geisser-Greenhouse correction revealed a significant main effect of delay (F (2.650, 29.15) = 25.92, p<0.0001), no main effect of drug treatment (F (1.000, 11.00) = 3.002, p=0.11) or virus (F (1, 11) = 1.893, p=0.19). Critically, we found a significant interaction effect (F (1, 11) = 8.045, p=0.016) of drug treatment (CNO vs SAL) and virus (hM4di vs control). There were no interactions between virus x delay (F (3, 33) = 1.748, p=0.18) nor drug treatment x delay (F (1.885, 20.74) = 1.032, p=0.37). Post-hoc analysis revealed a significant difference between CNO and saline treatment in the hM4Di group (F1,11=9.7, p=0.01), but not in the control group (F1,11=0.65, p=0.437) adjusted for multiple comparisons (Bonferroni) with each F testing the multivariate simple effects within each level. Since there were no interaction effects with delay, we combined all the delays and ran a two-way ANOVA mixed ANOVA with drug treatment (CNO vs SAL) as a within-subject factor and virus (hM4Di vs control) as a between-subject factor. We found a significant drug treatment x virus interaction effect, but no drug or virus main effect. Post-hoc test (Bonferroni correction for multiple comparisons) revealed a significant difference between SAL and CNO treatment in the rats that received the inhibitory DREADD virus (t=3.11, df=11, p=0.02), but not in rats that received the control virus (t=0.8, df=11, p=0.68).

We found that using an inhibitory chemogenetic approach targeting RE neurons that project to PrL did not alter DNMTP performance. We found no difference in the percent correct when rats received CNO in PrL in rats that received DREADD virus hM4Di and in rats that received the control virus in the RE. Specifically, a three-way mixed ANOVA with Geisser-Greenhouse correction revealed a significant main effect of delay (F (2.621, 41.93) = 31.16), no main effect of drug treatment (F (1.000, 16.00) = 0.32), virus (F (1, 16) = 0.01) and no interaction effects (drug treatment x virus: F (1, 16) = 1.4, virus x delay: F (3, 48) = 0.33, and drug treatment x delay: F (2.7, 43.14) = 0.76). We combined all the delays and ran a two-way ANOVA mixed ANOVA with drug treatment (CNO vs SAL) as a within-subject factor and virus (hM4di vs control) as a between-subject factor. We found no significant main effect of drug treatment (F (1, 16) = 0.32), virus (F (1, 16) = 0.01), and no drug treatment x virus interaction effect (F (1, 16) = 1.4).

## 4. Discussion

We found that pharmacological inactivation of the mPFC and RE impaired performance in the operant DNMTP in a delay-independent manner. Our finding that the mPFC is critical for working memory performance in Fischer 344 is consistent with previous work in Long-Evans rats showing a role of mPFC using both permanent lesions (Dunnett et al. 1999) and temporary GABA-mediated pharmacological inactivation (Auger and Floresco 2017), as well as improved working memory in aged Fischer rats with enhanced glutamatergic transmission in the mPFC (McQuail *et al*. 2016). Our findings that pharmacological inactivation of the RE impairs working memory is consistent with prior work in a spatial working memory delayed nonmatch task (T-maze) using optogenetic techniques (Maisson et al. 2018) but contrasts previous work with permanent lesions of RE showing improved performance in an operant DNMTP (Prasad et al. 2017). One possible reason behind this contradiction is a compensatory mechanism following the lesion leading to the improvement in working memory in the DNMTP task.

Using a chemogenetic approach to silence mPFC neurons that project to RE, we found that inhibiting the mPFC-RE circuit prior to DNTMP testing resulted in a delay-independent reduction in behavioral performance in experimental (hM4Di), but not control animals. We used a similar approach to determine if the reciprocal pathway (RE-mPFC) was also critical for working memory and found that manipulating the RE-mPFC did not impair performance. Our findings suggest a top-down (mPFC-RE) requirement for working memory as measured by DNMTP. In support, suppressing transmission in the mPFC-RE circuit using a chemogenetic approach abolished sequence memory which depends on functional working memory, while leaving general motivated behavior intact [i.e., speed, latency to retrieve rewards (Jayachandran *et al*. 2019)].

The RE is located in an advantageous anatomical position to relay information from the mPFC to the dorsal hippocampus that is required for memory encoding, given the lack of direct connection between the mPFC and dorsal hippocampus (Sesack *et al*. 1989; Buchanan *et al*. 1994). Thus, the RE is thought to play a key role in mediating prefrontal-hippocampal interactions to guide the ability to retain information and use it to guide memory. Indeed, there are dense bilateral projections between the mPFC, particularly the prelimbic (PrL) and infralimbic (IL) subregions, and the thalamic nuclei, RE (Vertes 2004). In support, stimulation of RE leads to elevated synchrony between the mPFC and hippocampus (Jayachandran et al. 2023), and inhibiting the RE leads to impairments in spatial working memory that depend on both the mPFC and hippocampus (Urban *et al*. 2014; Hallock et al. 2016; Maisson *et al*. 2018). The hippocampus is not necessary for performance in the operant version of the delayed (non)match to position task (Sloan *et al*. 2006), while the hippocampus is critical for delayed (non) matching performance in tasks that require spatial navigation such as a T-maze or water maze (Hampson *et al*. 1999; Nguyen *et al*. 2000; Zhang *et al*. 2013). Thus, our work extends on the previous findings in that the mPFC-RE is critical for working memory in the DNMTP task, which does not require hippocampus function (Sloan *et al*. 2006). While the RE is a critical interface between the mPFC and HPC for many types of memory functions including sequence learning (Jayachandran *et al*. 2019) and spatial working memory (Hallock *et al*. 2016; Maisson *et al*. 2018), our findings suggest a broader role of the mPFC-RE in working memory beyond the role of integrating the information with the hippocampus. In support, the inactivation of the mPFC leads to a decrease in burst firing in the RE *in vivo* (Zimmerman and Grace 2018) suggesting a functional role of mPFC on RE-dependent behavior.

RE cells that project to both mPFC and hippocampus are limited, but present with approximately 5-10% collateralization (Hoover and Vertes 2012; Varela et al. 2014). However, a greater percentage of RE cells that project to mPFC and the hippocampus are segregated, suggesting a potential separation of function. As such, the RE is located anatomically to modulate activity both a top-down and bottom-up functional approach (i.e., hippocampus-RE-mPFC and mPFC-RE-hippocampus). Since we did not find support for a role of RE-mPFC in the operant delayed nonmatch to position task, a more hippocampal-dependent task may be critical to engage the thalamic to mPFC circuitry. Indeed, different types of working memory engage the mPFC and the hippocampus differently (Kesner *et al*. 1996) with mPFC being more critical in visual-based working memory tasks, as such the RE may be an underlying link for working memory more generally.

Our findings, in conjunction with a significant amount of work examining the role of the mPFC-RE in sequence memory and spatial working memory, suggest that the mPFC-RE circuit plays a role in working memory function to maintain information online, regardless of the dependence of the hippocampus on the task. Thus, while the RE is critical for synchrony between the mPFC-HPC, and behavioral performance, RE function in the context of working memory goes beyond relaying information between mPFC and HPC. The mPFC plays a significant role in modulating expected outcome value to alter behavior in real-time across behavioral flexibility domains (Birrell and Brown 2000; Ostlund and Balleine 2005; Tran-Tu- Yen et al. 2009; West et al. 2021; Niedringhaus and West 2022), which is critical in a working memory task in which information needs to be held online trial by trial such as the DNMTP. RE is critical for behavioral flexibility in spatial navigation a win-stay task (Viena et al., 2018), as well as reversal learning and set-shifting (Rojas et al. 2024). As such, future studies can determine if the RE function extends beyond spatial behavioral flexibility and has a general role in the ability to maintain and manipulate behavior in real time across domains.

## Acknowledgments

This work was funded in part by the National Institute on Drug Abuse (R00DA042934 to EAW), the National Institute on Aging (R21AG072355 to EAW), and the National Institute of General Medical Sciences (T34GM127154 to TG), as well as Rowan University School of Osteopathic Medicine internal funds.

## Notes

### Competing Interest Statement

The authors have declared no competing interest.

## References

Auger ML, Floresco SB. 2017. Prefrontal cortical GABAergic and NMDA glutamatergic regulation of delayed responding. Neuropharmacology. 113:10–20.

Baddeley A. 1992. Working Memory: The Interface between Memory and Cognition. J Cogn Neurosci. 4:281–288.

Beas BS, Setlow B, Bizon JL. 2013. Distinct manifestations of executive dysfunction in aged rats. Neurobiol Aging. 34:2164–2174.

Benoit LJ, Holt ES, Teboul E, Taliaferro JP, Kellendonk C, Canetta S. 2020. Medial prefrontal lesions impair performance in an operant delayed nonmatch to sample working memory task. Behav Neurosci. 134:187–197.

Birrell JM, Brown VJ. 2000. Medial frontal cortex mediates perceptual attentional set shifting in the rat. J Neurosci. 20:4320–4324.

Buchanan SL, Thompson RH, Maxwell BL, Powell DA. 1994. Efferent connections of the medial prefrontal cortex in the rabbit. Exp Brain Res. 100:469–483.

Chen CM, Stanford AD, Mao X, Abi-Dargham A, Shungu DC, Lisanby SH, Schroeder CE, Kegeles LS. 2014. GABA level, gamma oscillation, and working memory performance in schizophrenia. Neuroimage Clin. 4:531–539.

Cho RY, Konecky RO, Carter CS. 2006. Impairments in frontal cortical gamma synchrony and cognitive control in schizophrenia. Proc Natl Acad Sci U S A. 103:19878–19883.

Churchwell JC, Morris AM, Musso ND, Kesner RP. 2010. Prefrontal and hippocampal contributions to encoding and retrieval of spatial memory. Neurobiol Learn Mem. 93:415–421.

Cohen RM, Rezai-Zadeh K, Weitz TM, Rentsendorj A, Gate D, Spivak I, Bholat Y, Vasilevko V, Glabe CG, Breunig JJ, Rakic P, Davtyan H, Agadjanyan MG, Kepe V, Barrio JR, Bannykh S, Szekely CA, Pechnick RN, Town T. 2013. A transgenic Alzheimer rat with plaques, tau pathology, behavioral impairment, oligomeric abeta, and frank neuronal loss. J Neurosci. 33:6245–6256.

Dolleman-van der Weel MJ, Griffin AL, Ito HT, Shapiro ML, Witter MP, Vertes RP, Allen TA. 2019. The nucleus reuniens of the thalamus sits at the nexus of a hippocampus and medial prefrontal cortex circuit enabling memory and behavior. Learn Mem. 26:191–205.

Dunnett SB, Nathwani F, Brasted PJ. 1999. Medial prefrontal and neostriatal lesions disrupt performance in an operant delayed alternation task in rats. Behav Brain Res. 106:13–28.

Euston DR, Gruber AJ, McNaughton BL. 2012. The role of medial prefrontal cortex in memory and decision making. Neuron. 76:1057–1070.

Ferretti MT, Iulita MF, Cavedo E, Chiesa PA, Schumacher Dimech A, Santuccione Chadha A, Baracchi F, Girouard H, Misoch S, Giacobini E, Depypere H, Hampel H, Women’s Brain P, the Alzheimer Precision Medicine I. 2018. Sex differences in Alzheimer disease - the gateway to precision medicine. Nat Rev Neurol. 14:457-469.

Gohar T, Ciacciarelli EJ, Dunn SD, West EA. 2023. Transient strain differences in an operant delayed non-match to position task. Behav Processes. 211:104932.

Guarino A, Favieri F, Boncompagni I, Agostini F, Cantone M, Casagrande M. 2018. Executive Functions in Alzheimer Disease: A Systematic Review. Front Aging Neurosci. 10:437.

Haenschel C, Bittner RA, Waltz J, Haertling F, Wibral M, Singer W, Linden DE, Rodriguez E. 2009. Cortical oscillatory activity is critical for working memory as revealed by deficits in early-onset schizophrenia. J Neurosci. 29:9481–9489.

Hallock HL, Wang A, Griffin AL. 2016. Ventral Midline Thalamus Is Critical for Hippocampal-Prefrontal Synchrony and Spatial Working Memory. J Neurosci. 36:8372–8389.

Hampson RE, Jarrard LE, Deadwyler SA. 1999. Effects of ibotenate hippocampal and extrahippocampal destruction on delayed-match and -nonmatch-to-sample behavior in rats. J Neurosci. 19:1492–1507.

Hoover WB, Vertes RP. 2012. Collateral projections from nucleus reuniens of thalamus to hippocampus and medial prefrontal cortex in the rat: a single and double retrograde fluorescent labeling study. Brain Struct Funct. 217:191–209.

Jayachandran M, Linley SB, Schlecht M, Mahler SV, Vertes RP, Allen TA. 2019. Prefrontal Pathways Provide Top-Down Control of Memory for Sequences of Events. Cell Reports. 28:640–654.e646.

Jayachandran M, Viena TD, Garcia A, Veliz AV, Leyva S, Roldan V, Vertes RP, Allen TA. 2023. Nucleus reuniens transiently synchronizes memory networks at beta frequencies. Nat Commun. 14:4326.

Kesner RP, Hunt ME, Williams JM, Long JM. 1996. Prefrontal cortex and working memory for spatial response, spatial location, and visual object information in the rat. Cereb Cortex. 6:311–318.

Kirova AM, Bays RB, Lagalwar S. 2015. Working memory and executive function decline across normal aging, mild cognitive impairment, and Alzheimer’s disease. Biomed Res Int. 2015:748212.

Kumar S, Zomorrodi R, Ghazala Z, Goodman MS, Blumberger DM, Cheam A, Fischer C, Daskalakis ZJ, Mulsant BH, Pollock BG, Rajji TK. 2017. Extent of Dorsolateral Prefrontal Cortex Plasticity and Its Association With Working Memory in Patients With Alzheimer Disease. JAMA Psychiatry. 74:1266–1274.

Maisson DJN, Gemzik ZM, Griffin AL. 2018. Optogenetic suppression of the nucleus reuniens selectively impairs encoding during spatial working memory. Neurobiology of Learning and Memory. 155:78–85.

McQuail JA, Beas BS, Kelly KB, Simpson KL, Frazier CJ, Setlow B, Bizon JL. 2016. NR2A- Containing NMDARs in the Prefrontal Cortex Are Required for Working Memory and Associated with Age-Related Cognitive Decline. J Neurosci. 36:12537–12548.

Mielke MM. 2018. Sex and Gender Differences in Alzheimer’s Disease Dementia. Psychiatr Times. 35:14–17.

Nguyen PV, Abel T, Kandel ER, Bourtchouladze R. 2000. Strain-dependent differences in LTP and hippocampus-dependent memory in inbred mice. Learn Mem. 7:170–179.

Niedringhaus M, West EA. 2022. Prelimbic cortex neural encoding dynamically tracks expected outcome value. Physiol Behav. 256:113938.

Ostlund SB, Balleine BW. 2005. Lesions of medial prefrontal cortex disrupt the acquisition but not the expression of goal-directed learning. J Neurosci. 25:7763–7770.

Paxinos G, Watson C. 2013. The Rat Brain in Stereotaxic Coordinates: Elsevier Science.

Prasad JA, Abela AR, Chudasama Y. 2017. Midline thalamic reuniens lesions improve executive behaviors. Neuroscience. 345:77–88.

Rojas AKP, Linley SB, Vertes RP. 2024. Chemogenetic inactivation of the nucleus reuniens and its projections to the orbital cortex produce deficits on discrete measures of behavioral flexibility in the attentional set-shifting task. Behav Brain Res. 470:115066.

Rorabaugh JM, Chalermpalanupap T, Botz-Zapp CA, Fu VM, Lembeck NA, Cohen RM, Weinshenker D. 2017. Chemogenetic locus coeruleus activation restores reversal learning in a rat model of Alzheimer’s disease. Brain. 140:3023–3038.

Salat DH, Kaye JA, Janowsky JS. 2001. Selective preservation and degeneration within the prefrontal cortex in aging and Alzheimer disease. Arch Neurol. 58:1403–1408.

Sesack SR, Deutch AY, Roth RH, Bunney BS. 1989. Topographical organization of the efferent projections of the medial prefrontal cortex in the rat: an anterograde tract-tracing study with Phaseolus vulgaris leucoagglutinin. J Comp Neurol. 290:213–242.

Sloan HL, Good M, Dunnett SB. 2006. Double dissociation between hippocampal and prefrontal lesions on an operant delayed matching task and a water maze reference memory task. Behavioural Brain Research. 171:116–126.

Tran-Tu-Yen DA, Marchand AR, Pape JR, Di Scala G, Coutureau E. 2009. Transient role of the rat prelimbic cortex in goal-directed behaviour. Eur J Neurosci. 30:464–471.

Urban KR, Layfield DM, Griffin AL. 2014. Transient inactivation of the medial prefrontal cortex impairs performance on a working memory-dependent conditional discrimination task. Behav Neurosci. 128:639–643.

Varela C, Kumar S, Yang JY, Wilson MA. 2014. Anatomical substrates for direct interactions between hippocampus, medial prefrontal cortex, and the thalamic nucleus reuniens. Brain Struct Funct. 219:911–929.

Vertes RP. 2002. Analysis of projections from the medial prefrontal cortex to the thalamus in the rat, with emphasis on nucleus reuniens. J Comp Neurol. 442:163–187.

Vertes RP. 2004. Differential projections of the infralimbic and prelimbic cortex in the rat. Synapse. 51:32–58.

Webb AA, Gowribai K, Muir GD. 2003. Fischer (F-344) rats have different morphology, sensorimotor and locomotor abilities compared to Lewis, Long-Evans, Sprague-Dawley and Wistar rats. Behav Brain Res. 144:143–156.

West EA, Forcelli PA, Murnen AT, McCue DL, Gale K, Malkova L. 2012. Transient inactivation of basolateral amygdala during selective satiation disrupts reinforcer devaluation in rats. Behav Neurosci. 126:563–574.

West EA, Niedringhaus M, Ortega HK, Haake RM, Frohlich F, Carelli RM. 2021. Noninvasive Brain Stimulation Rescues Cocaine-Induced Prefrontal Hypoactivity and Restores Flexible Behavior. Biol Psychiatry. 89:1001–1011.

Wicker E, Hyder SK, Forcelli PA. 2022. Pathway-specific inhibition of critical projections from the mediodorsal thalamus to the frontal cortex controls kindled seizures. Prog Neurobiol. 214:102286.

Zhang XH, Liu SS, Yi F, Zhuo M, Li BM. 2013. Delay-dependent impairment of spatial working memory with inhibition of NR2B-containing NMDA receptors in hippocampal CA1 region of rats. Mol Brain. 6:13.

Zimmerman EC, Grace AA. 2018. Prefrontal cortex modulates firing pattern in the nucleus reuniens of the midline thalamus via distinct corticothalamic pathways. Eur J Neurosci. 48:3255–3272.

